# Integrating Rigidity Analysis into the Exploration of Protein Conformational Pathways using RRT* and MC

**DOI:** 10.1101/2021.04.09.439189

**Authors:** Fatemeh Afrasiabi, Ramin Dehghanpoor, Nurit Haspel

## Abstract

To understand how proteins function on a cellular level, it is of paramount importance to understand their structures and dynamics, including the conformational changes they undergo to carry out their function. For the aforementioned reasons, the study of large conformational changes in proteins has been an interest to researchers for years. However, since some proteins experience rapid and transient conformational changes, it is hard to experimentally capture the intermediate structures. Additionally, computational brute force methods are computationally intractable, which makes it impossible to find these pathways which require a search in a high-dimensional, complex space. In our previous work, we implemented a hybrid algorithm that combines Monte-Carlo (MC) sampling and RRT*, a version of the Rapidly Exploring Random Trees (RRT) robotics-based method, to make the conformational exploration more accurate and efficient, and produce smooth conformational pathways. In this work, we integrated the rigidity analysis of proteins into our algorithm to guide the search to explore flexible regions. We demonstrate that rigidity analysis dramatically reduces the run time and accelerates convergence.

## 1. Introduction

Proteins take part in nearly all biological functions in the cell. For this reason, it is of paramount importance to understand how they function. A protein’s biological function is dictated by its 3-dimensional structure and its interaction with other molecules. From the structures of proteins, we have derived our understanding both of the functions of individual proteins, and the general principles of protein structure and dynamics. Proteins fold into unique 3D shapes, and as they are dynamic molecules, they may change their shapes as a consequence of environmental changes or binding to other molecules. The rearrangement of a protein’s tertiary structure in response to environmental changes, external factors, or binding of a ligand from one conformation to another is called the conformational change. For these reasons, in order to fully elucidate the functionality of proteins, characterizing these conformational changes, which are significant factors in protein properties, has been a continuing hotspot for research [1–4].

However, one of the main challenges is that these conformational changes are transitory and that often makes it hard to characterize the intermediate structures. Researchers have used computational methods to try to capture these computational changes and the paths the proteins take when they transition from one conformation to another. For instance, Molecular Dynamics (MD) simulations [5] can examine protein dynamics up to the microsecond level though they are still not fast enough to capture conformational transitions that take place over larger timescales - milliseconds or more. Methods such as targeted, biased, or steered MD, umbrella sampling, replica exchange, activation relaxation, conformational flooding, swarm methods, and others attempt to lower the computational demands of MD-based approaches [6–9], and Multiscale Molecular Dynamics Simulation methods [10]. Some methods focus on deforming a trivial conformational path (obtained, for instance, through morphing) to improve its energy profile. MD-based approaches tend to produce one path, depending on the starting conditions. Another class of methods samples the conformational space efficiently by perturbing the molecular structure at certain degrees of freedom and using geometric and biophysical constraints to guide the search. The search is done using various techniques such as normal mode analysis (NMA) [11,12], elastic network [13,14], robotic motion [15] planning, and more. These methods are usually faster than MD but do not result in physical pathways due to randomness in the search and the approximations being used [16]. Therefore, further processing should often be applied to extract biologically meaningful information. Several motion-planning methods were developed in the past to sample the conformational space of proteins using prior knowledge [17] or multi-scale sampling [18–20]. A brief review of adaptive sampling methods is in [21]. Efficient and accurate sampling of protein conformational pathways helps to better identify intermediate states which also correspond to local energy minima. The SPIRAL method described in [22] tries to solve the same problem of simulating molecular motion, but is computationally expensive. A novel evolutionary algorithm that aims to sample regions of interest in the conformational space is presented in [23]. It presents a promising research direction in improving decoy generation for template-free protein structure prediction. A similar approach is proposed in [24] by using a highly parallelized version of RRT called Transition-based RRT. This algorithm also aims at identifying the most probable conformations of a class of highly-flexible biomolecules such as Intrinsically Disordered Proteins (IDPs).

In [18], the authors combined methods originating from robotics and computational biophysics, to model protein conformational transitions. To reduce the dimension of the problem, normal modes are computed for a coarse-grained elastic network model built on short fragments of three residues. This added bias helps to efficiently sample conformational search space. While the incorporation of a suitable bias towards the goal has an impact on the accuracy of the simulation, structure forces reduce dwell time in a given stable or meta-stable state, the bias possibly sacrifices a more expansive view of possibly different transition trajectories to the goal structure. This is typically addressed by repeating the simulation to obtain many transition trajectories, which taken together can cover the transition ensemble in the absence of correlations between trajectories. A brief review of the current methods and their comparison can be found in [25].

A different class of methods focus on computing a sequence of conformations (a conformational path) with a credible energy profile. The working assumption is that credible conformational paths can then be locally deformed with techniques that consider dynamics to obtain transition trajectories. Methods in this category adapt sampling-based search algorithms [26] developed for the robot motion-planning problem which bears strong analogies to the problem of computing conformations along with a structural transition. There are some unique challenges offered up by molecular systems. First, molecular systems typically have an exceptionally high number of degrees of freedom. Second, the energy surface of a single protein is highly complex with many local minima. The methods developed in algorithmic robotics to address the robot motion planning problem fall under either the roadmap-based or tree-based category. Roadmap-based methods [22] are limited by the *steering problem*, which means, if one has two sampled conformations, it may not be possible to find a constraint-satisfying path steering the system from one conformation to another. This method uses a low-dimensional projection of the sampled conformations to approximate coverage of the conformational space.

### 1.1. RRT, RRT*, and Adaptive Sampling

The Rapidly Exploring Random Tree (RRT) algorithm [27] is a robotics-based method that builds a tree rooted at the start configuration by randomly exploring the search space. The algorithm was presented in our previous work, extended by this journal submission [28]. The algorithm is outlined here for clarity.

- Create a random new sample *q_r_* (a new molecular conformation).
- Find *q_r_*’s nearest neighbor (*q_near_*) on the tree.
- Try to connect *q_r_* and *q_near_* by an edge. To do this, we incrementally rotate the neighbor’s relevant backbone dihedral angles in the direction of the new conformation.
- Stop when you reach an obstacle (a high energy barrier) or when you reach the newly created sample. The stopping point is *q_new_*
- Add *q_new_* to the tree with *q_near_* as its parent.
- Stop when you reach an RMSD threshold distance from the goal or when the maximum number of iterations was reached.

RRT is known for its ability to rapidly explore the work space. However, due to its randomness, it expands to all areas of the search space, which means that it does not always converge fast, especially in high-dimensional search spaces like the protein conformational landscape. It also makes the resulting paths from start to goal jagged and “zigzagged”.

RRT* [29] is an optimized version of RRT. Due to the random nature of the original RRT’s node selection and connection, it does not always produce smooth or optimized paths. RRT* modifies the original algorithm to deliver significantly less ragged and smoother paths to the goal. The optimization is done through a heuristic cost function as follows:

- For every node we also record the tree distance from the root. After finding the nearest neighbor on the tree, we examine the neighborhood of the new node in a fixed radius. If there is a node with a cheaper overall cost, it becomes the node’s parent instead of the original nearest neighbor.
- After adding the new node to its cheapest neighbor, the paths are rewired: The neighbors are checked if being rewired to the newly added vertex will decrease their overall cost.
- If the cost indeed decreases, the neighbor is rewired to the newly added vertex. This step makes the path smoother and less jagged looking.

Figure 1 illustrates the process.

**Figure 1.**
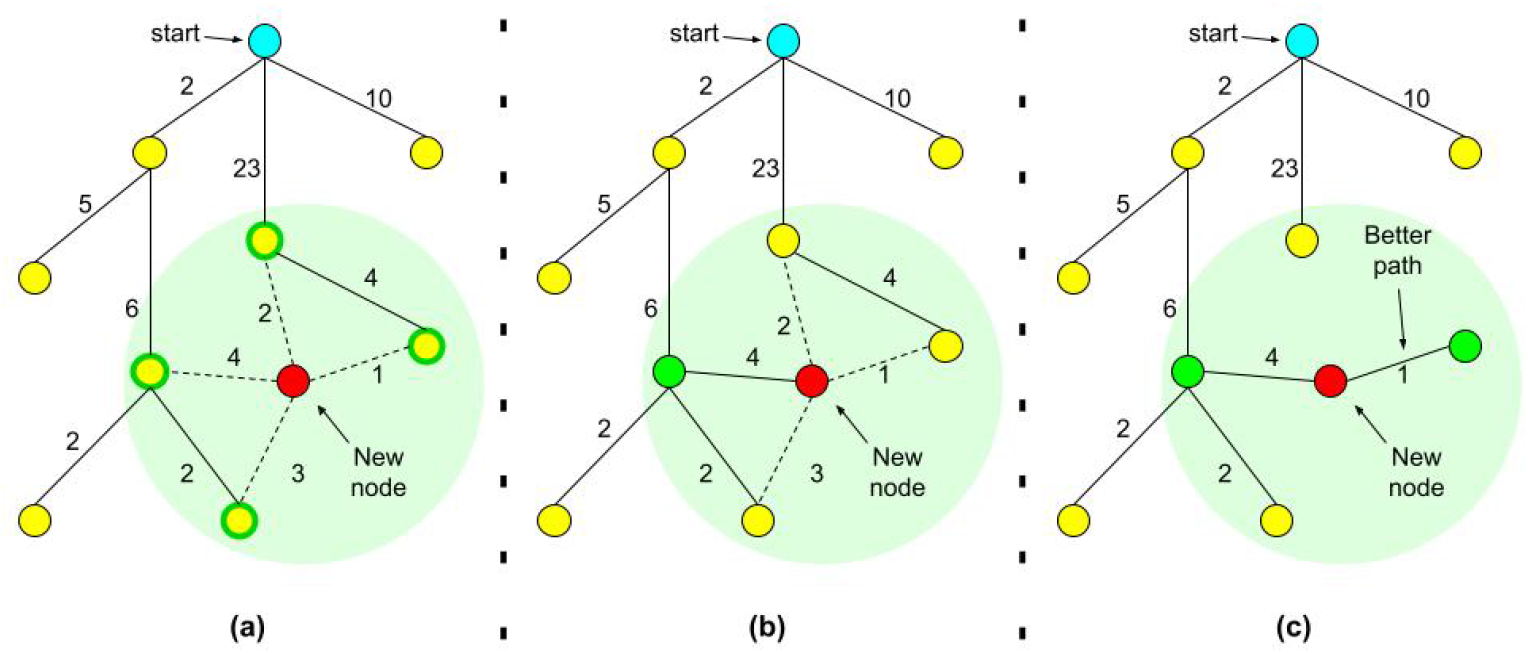
Illustration of RRT* best (least-cost) parent search and the rewiring process. (**a**) A new node is added to the tree and nearest neighbors are highlighted. (**b**) The best parent is chosen among nearest neighbors. (**c**) Costs of rewiring are checked and the tree is rewired.

### 1.2. Rigidity Analysis

The analysis of rigid residues in proteins can help predict which groups of atoms are more likely to move together in a coordinated way (which are called rigid clusters), and need more energy in order to be destabilized and deformed. These rigid areas of proteins are more likely to stay together during the conformational changes of proteins, since structural changes tend to happen around flexible hinges in the protein. Rigidity analysis methods [30,31] use only inter-atomic connectivity and interaction information and do not rely on energy calculations. In our work, we used the Kinari software [30,32] where the rigidity analysis of a protein is done in the following way: During data input and curation (first phase), a PDB file is retrieved from the Protein Data Bank site or uploaded by the user, chains and atoms to be retained are selected, and chemical interactions are calculated and can be removed, added, or modified. During the second phase, modeling options are specified, rigidity analysis is performed, and the results are explored with a script-enhanced Jmol-based (http://www.jmol.org) 3D visualizer.

The Kinari software is written in C++, and its library (KINARI-Lib) implements the pebble game algorithm. It also provides support for bar-joint and body-bar-hinge mechanical models. Pebble game rigidity analysis is an efficient method for extracting rigidity and flexibility information of biomolecules without having to perform costly molecular dynamics simulations [33]. This algorithm works on a multigraph associated with a mechanical model that is constructed from an arbitrary atom-bond network. In the Kinari software, the authors have developed a faster and more robust variation of the pebble game algorithm tailored to the specificities of biopolymers.

### 1.3. Our Previous Work

In our previous work we implemented a biased Monte Carlo tree search, to explore the conformational space of medium and large proteins [28,34]. In order to decrease the search time and at the same time increase the convergence of the search, we added an optional input argument to allow the algorithm to use a list of residues to manipulate during the search. These residues are assumed to either be co-evolving or otherwise important to the protein structure, function, or dynamics. If no such information is given, any residue may be selected for modification.

At the next step, our goal was to enhance the convergence time even more, as it was still time-consuming for large proteins, and also to generate smooth, even, and least-cost pathways. To do so, we incorporated the RRT* algorithm, the improved and optimized version of the robotics-based Rapidly Exploring Random Tree algorithm, and introduced three key methods, **nearNeighbors**, **chooseParent**, and **rewire** [28]. The **nearNeighbors** method will look for a set of nearest neighbors that lie within an RMSD of at most 1Å from each anew generated node. Then comes the **chooseParent** method that selects the least-cost node amongst the near neighbors to be the parent of the newly created node. We used an A*-based heuristic cost function to determine the cost of each node (neighbor) and choose the least-cost one. Then the **rewire** method was called to check if we can refine the paths we already had in the tree and make them more straight, as was the goal in RRT*’s creation for robot motion planning [29]. These methods helped the algorithm to generate smoother and less jagged pathways with minimum accumulated costs. In the end, when the tree expands beyond a certain size (dependent on the number of residues), the algorithm prunes the tree by removing the nodes that are within a threshold distance to one another to decrease redundancy.

### 1.4. This Contribution

This contribution is an extension of our conference paper described in [28]. Instead of using an optional list of residues for manipulation, our goal is to focus explicitly on flexible residues. In this work we use *rigidity analysis* to steer and guide the algorithm towards manipulating flexible hinges that are more likely to undergo large-scale changes. Rigidity analysis represents the protein as a set of rigid clusters, connected by flexible residues. This can assist the algorithm to reduce the randomness of choosing dihedral angles to perturb, and instead, focus on the flexible residues of the protein and select those for perturbing and creating new conformations. Since the proteins undergo large-scale conformational changes, their rigidity properties may change too. Therefore, we run rigidity analysis several times during the search and modify our criteria for selecting residues to manipulate.

## 2. Materials and Methods

### 2.1. Description of the algorithm

The input to the algorithm is two PDB coordinates representing conformations of the same protein, denoted as start and end conformations. The goal is to start with the source node, which represents the start conformation, and get as close to the target node, which we denote the end conformation. The algorithm produces pathways made of successive conformations, simulating the path that the protein takes when transitioning between the conformations. The algorithm continues to generate conformations until it reaches a conformation whose RMSD to the end conformation is less than a pre-determined threshold or until a maximum number of iterations is reached. The path is returned as output. An additional input source that we focus on in this work is a list of flexible residues. The list of flexible residues generated by the Kinari software [32] (details mentioned in the next subsection) is used when the algorithm wants to select certain bonds for manipulation. We use semi coarse-grained protein structural representations. Proteins are represented using their backbone chain and C-*β* atoms (except for Gly, which does not have a C-*β* atom). The degrees of freedom for each protein molecule are the *ϕ* and *ψ* angles around the C-*α* atoms. This way of representation of the proteins hits a balance between accuracy and speed efficiency.

#### The algorithm description follows

During each iteration, a conformation is chosen from the RRT tree to be perturbed. The tree at the beginning only contains the start conformation. We use a biased variable so that one-third of the time the conformation to be perturbed is selected out of the *k* best conformations. By best, we mean the closest, in terms of RMSD, to the end conformation. In this work, we chose *k* = 20. The rest of the time the conformation is selected from the general pool of nodes, which has all the nodes that are possible to create by manipulating the dihedral angles of molecular structure. The general pool of nodes represents the sampled conformations so far. Once a node is selected for expansion, the algorithm perturbs the molecular conformation of this node at numerous, randomly chosen, *ϕ* and *ψ* angles in order to generate a new conformation. These angles are randomly chosen from all the angles of the selected conformation but with the constraint that each particular angle’s difference from its counterpart in the end, aka target, conformation is larger than a certain threshold (In this work threshold = 5 degrees). Generally, these angles are selected from all the residues of the conformation, and that’s what we did in our previous work. But in this work, we set aside the rigid residues of the conformation and instead give the algorithm a list of flexible residues to choose these angles from. Since these are the residues that are not part of any rigid clusters, they are more likely to be different in the target conformation and hence are good candidates to be perturbed. After generating the new random conformation by perturbing the angles of flexible residues, it undergoes *k* steps of steepest descent energy minimization in order to relax local clashes while not altering the structure of the conformation a lot. The constants that we use in this work are *k* = 10 for the steps of steepest descent, and 5 * *thenumberof residues* as a threshold for acceptable energy. If the new conformation has an energy value above the acceptable threshold after the steps of steepest descent, it is accepted as a new node to our pool of conformations based on a Monte-Carlo criterion. Next, in the nearNeighbors method, the algorithm looks for a set of nearest neighbors (as the RRT* algorithm suggests to find neighbors that exist at a certain distance from the new node) which in our case are the nodes that exist within an RMSD of at most 1 Å from the newly added node. In case there are no other nodes within this RMSD, the nearest existing node would be the only node selected to create this set of nearest neighbors. Among these nearest neighbors, the best (defined by our cost method) least-cost node is selected to be set as the parent of this new node in the tree. The cost of each node is calculated using an A*-based heuristic cost function. The cost function used here is

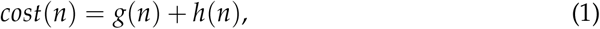

where *g*(*n*) is the summation of the *g*(*parent*(*n*)) and the RMSD of *n* from the start conformation, and *h*(*n*) is the RMSD of *n* from the end conformation. We implemented this cost function to estimate the cost of the cheapest (least-cost) path from the newly generated conformation to the end conformation. The least-cost node is chosen in the chooseParent method and is set as the parent of the new node and added to the main tree. The tree goes through a rewiring procedure afterward, where we check if the newly added node to the tree is a better parent for its neighbors than their current ones. That means, for each node in the set of nearest neighbors, we check if the cost to get that node is less through our newly added node than its current parent. And if so, we replace the current parent with the new node.

After multiple iterations, the pool of conformations that we have tends to get larger and larger and it becomes both space and time-wasting to look through all these conformations. Therefore, we go through the pool to find “redundant” nodes, and by that we mean the nodes whose RMSD from each other is 10% or less than their RMSD from the end conformation, and only keep the one closer to the end conformation. This pruning process happens first when the size of the pool tree is equal to the protein molecule’s number of residues and then every time it is multiplied by a constant (we chose 3 experimentally). The potential energy function that we use to calculate the energy of a new conformation, in order to decide whether to accept it or not is as follows: [35]

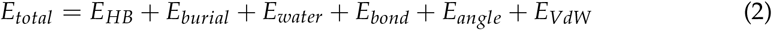

The first three terms represent hydrogen bonds, burial and water mediated interaction terms taken from [35]. The three remaining terms *E_VdW_, E_bond_*, and *E_angle_* came from the *AMBER f f* 03 force field [36]. The *E_VdW_* is a Lennard Jones potential that was slightly adapted to tolerate soft collisions between atoms, as a result of the lower resolution. These terms were used to test for clashes and deformities in the structure. A new molecule’s potential energy must be smaller than a particular threshold, i.e., *k*\* the number of residues in the molecule, to be accepted to the pool. In this paper we set *k* = 5. If the energy was too high, we attempted to minimize it using up to 10 steps of gradient-descent minimization on the bond, angle and VdW terms. If the minimization failed to bring the conformation to within the threshold, it was discarded. If the potential energy was below the threshold, the conformation was accepted to be added to the tree.

The new conformation is now accepted and the next step is to make a new branch (path of conformations) towards it. We do this considering the below criteria:

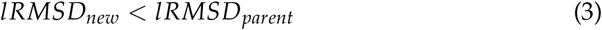

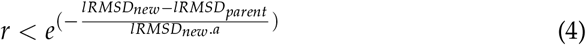

where *r* is a random number in [0,1), and *a* is a previously-defined constant. *a* is defined as 0.01 in this work. To avoid getting stuck in the potential local minima, with a small possibility, we allow conformations with lower scores to be accepted.

The iterative sampling of new conformations proceeds until either the score of the generated conformation is below the preset score threshold or the minimum score achieved by the conformations in the pool does not reduce for *M* = 500 iterations. We measure the score of each conformation by its *lRMSD* with respect to the goal conformation, and the threshold value that we set for this score is within 1.7 and 3.65 based on the protein size and the difficulty of the pathway. This range is a realistic lower bound we found for our tested proteins since achieving a lower *lRMSD* is challenging because of sampling errors and variations of the protein molecules The very last step is to extract a path from the tree, from the start conformation to the goal conformation when the threshold *lRMSD* is reached.

### 2.2. Integrating Rigidity Analysis

The highlight of this contribution is integrating the rigidity analysis of the protein into our algorithm to steer the algorithm when it selects dihedral angles to manipulate during the search. The information regarding rigidity analysis can be gathered from computational analysis [37,38], experimental information, co-evolution, or rigidity analysis tools such as Kinari [30,32]. For each protein, We used Kinari [30] to obtain the rigid clusters of residues and then extract the flexible residues to be manipulated by our algorithm. The output from Kinari contains the atoms that belong to rigid clusters of any size. In order to avoid very small rigid clusters which may contain noise, we set a threshold for the cluster sizes to be considered. We experimented with thresholds of 6, 7, and 8 atoms. Any residue with atoms not in a rigid cluster was considered flexible. We investigate the effectiveness of using rigidity analysis to guide the residue-selecting procedure in two ways. One approach is to incorporate rigidity analysis only at the beginning of the algorithm as described below:

- We generate the start conformation’s rigid clusters of atoms, using a specified threshold for the minimum number of atoms per rigid cluster.
- The flexible residues are then produced using the rigid clusters extracted from the previous step.
- The list of flexible residues of the start conformation is given to the algorithm to be used as candidate residues for perturbation of their *ϕ* and *ψ* dihedral angles.

In the above description rigidity analysis is only run once in the beginning, but since the protein undergoes large conformational changes, its rigid clusters may change during the run. The second approach is to use rigidity analysis not only at the beginning but also throughout the algorithm to account for these changes. We hypothesized that feeding the algorithm with the results of rigidity analysis would make the algorithm produce smoother pathways and eventually converge faster. We found that the optimal number of rigidity analysis recalculations was when the conformation changes by more than 2Å after the previous run. This results in approximately 2–7 runs in the test cases studied here.

### 2.3. Using The Kinari Software

The Kinari software gets the PDB ID and the chain ID of a protein and returns the set of rigid clusters of atoms. Each rigid cluster is shown as a list of atom numbers. Small clusters with very few atoms (defined by a lower limit) are disregarded because they might represent for example a Benzene ring whose rigidity is of no interest because of the inherent planarity of such structures. We used three different values as the lower limit of atoms in a rigid cluster, 6, 7, and 8, and ran our algorithm for all of these values to see whether this lower limit has a genuine impact on the convergence or the runtime of the algorithm. For instance, by choosing 6 as the lower limit, all of the rigid clusters discovered with fewer than six atoms were not considered.

### 2.4. Implementation Details

In our previous work, we tested our algorithm on different proteins by submitting jobs on UMass Boston compute cluster, Chimera. Chimera is a heterogeneous distributed memory high-performance compute cluster. However, we were incapable of running the Kinari software on Chimera by submitting jobs through the job scheduler and had to run the algorithm on a new operating system on Chimera without using the job scheduler. Because of this change, we re-ran our previous algorithm [28] on our test cases on the same new OS as the current implementation to be able to compare the results and runtimes properly. Each test case was run ten times and the results were averaged. We used the same RMSD threshold for all runs, and the RMSD results were approximately the same per protein in all implementations regardless of rigidity analysis. Table 1 shows the comparison of three versions of our algorithm. And table 2 estimates the memory usage by comparing the resulting tree size.

**Table 1.** Performance and Comparison of the Results.

| Molecule | Path | Init. RMSD<sup>†</sup> | min RMSD | Time<sup>a</sup> | Time<sup>b</sup> | Time<sup>c</sup> |
| --- | --- | --- | --- | --- | --- | --- |
| CaM | 1CLL → 1CTR | 14.85 | 2.94 | 92.20 | 118.2 | 63.02 |
|  | 1CTR → 1CLL | 14.85 | 2.93 | 59.75 | 67.43 | 29.19 |
|  | 2F3Y → 1CFD | 9.93 | 3 | 58.67 | 49.01 | 37.80 |
|  | 1CFD → 2F3Y | 9.93 | 3.57 | 45 | 40.38 | 32 |
| AdK | 1AKE → 4AKE | 7.14 | 2.59 | 411.67 | 212.2 | 182.73 |
|  | 4AKE → 1AKE | 7.14 | 2.68 | 307 | 186.75 | 142.69 |
| CVN | 1L5E → 2EZM | 15.92 | 2.93 | 73.5 | 68.4 | 56 |
|  | 2EZM → 1L5E | 15.92 | 2.87 | 33 | 32.67 | 29.82 |
| RBP | 2DRI → 1URP | 4.08 | 2.37 | 26.33 | 6.67 | 4.33 |
|  | 1URP → 2DRI | 4.08 | 2.42 | 23 | 5.75 | 4.33 |

<sup>†</sup> The distances are in Å and the time in minutes.
<sup>a</sup> Runtime without rigidity analysis (our previous algorithm)
<sup>b</sup> Runtime with rigidity analysis of the start conformation only at the beginning
<sup>c</sup> Runtime with rigidity analysis throughout the algorithm

**Table 2.** Comparison of memory usage, size of the trees created

| Molecule | Path | Tree size <sup>† a</sup> | Tree size <sup>b</sup> | Tree size <sup>c</sup> |
| --- | --- | --- | --- | --- |
| CaM | 1C$\text{LL}$ → 1C$\text{TR}$ | 2209 | 2700 | 2137 |
| | 1C$\text{TR}$ → 1C$\text{LL}$ | 1597 | 1708 | 1166 |
| | 2F3Y → 1CFD | 1738 | 1603 | 1488 |
| | 1CFD → 2F3Y | 1432 | 1410 | 1354 |
| AdK | 1AKE → 4AKE | 2398 | 1661 | 1829 |
| | 4AKE → 1AKE | 1526 | 1491 | 1473 |
| Cvn | 1L5E → 2EZM | 3279 | 2886 | 3059 |
| | 2EZM → 1L5E | 1921 | 1680 | 1867 |
| Rbp | 2DRI → 1URP | 100 | 91 | 84 |
| | 1URP → 2DRI | 109 | 84 | 63 |

<sup>†</sup> Number of nodes in the RRT tree generated
*a* without rigidity analysis (our previous algorithm)
*b* with rigidity analysis of the start conformation only at the beginning
*c* with rigidity analysis throughout the algorithm

## 3. Results

Below we detail the test case we used and we elaborate on the results for each system separately. Table 1 shows the comparison of three versions of our algorithm: One denoted by (a) for our previous work [28], one denoted by (b) for the version that runs rigidity analysis once in the beginning and one, denoted (c), which runs rigidity analysis throughout the algorithm (see Methods for details). In all cases, the run time improved considerably in implementation (c) with respect to (a). In all but Calmodulin implementation (b), which performed one rigidity analysis, improved the run time considerably. The memory consumption, measured in the size of the tree, also improved when running rigidity analysis, but the difference is less marked, possibly due to the tree pruning procedures discussed above. Below we detail the results for each protein.

### 3.1. Test Cases

#### Calmodulin (CaM)

Calmodulin is a highly conserved, calcium-modulated protein that is found in all eukaryotic cells. It is the most adaptable member of its family, the family of calcium-binding proteins. It acts as a receptor for the *Ca*^2+^ signal [39]. Calmodulin modulates many cellular effects such as regulation of changes in the concentration of cytosolic calcium ions that control biological processes [40]. It is a relatively small protein consisting of 148 amino acids. Calmodulin has two globular domains that are relatively symmetrical. Each of these domains contains a pair of EF-hand motifs^1^, and the N- and C-domain motifs that are separated by a flexible linker region. This would make a total of four *Ca*^2+^ binding sites in this protein.

We considered two Calmodulin pathways for our simulations. First, we simulated the pathway between an open conformation (PDB: 1*CLL*) and a closed conformation (PDB:1*CTR*) The RMSD between the two conformations is 14.83Å. We ran our simulations in both directions, from the open conformation to the closed conformation and then the other way around. Figure 2 (a) and (b) show the closed (1*CTR*) and open (1*CLL*) conformations, respectively. And Figure 3 shows the superimposition of the simulated conformation and the actual conformation of the protein molecule. The flexible residues detected by rigidity analysis are also represented in Figure 2 in blue. As expected, the flexible residues are predicted to be mostly on the linker region between the two EF-hands of the protein. The predicted flexible residues are not exactly the same as the real changed residues when the algorithm gets to the end conformation, but they mostly cover the main flexible parts and guide the algorithm to converge faster towards the end conformation. Table 1 shows a comparison of the results of from our previous work [28], the version that uses the flexible (non-rigid) residues of the start conformation throughout the run, and our current works that uses rigidity analysis to create flexible residues several times throughout the simulation as explained in Section 2.2. For this pathway, when the algorithm calls the Kinari software several times throughout the run, we get the fastest convergence. For instance, for 1*CTR* → 1*CLL* our runtime average is 30.56 minutes faster than our previous work (Table 1), and for 1*CLL* → 1*CTR*, the new runtime is 29.18 minutes faster. Table 2 displays the size of the RRT tree for our three scenarios. By comparing columns a and c, we can see in all cases, the size of the tree is reduced when we incorporate rigidity analysis a couple of times throughout the algorithm. This denotes a decrease in memory usage. For 1*CTR* → 1*CLL*, RRT tree size changed from 1597 to 1166 and for 1*CLL* → 1*CTR*, RRT tree size changed from 2209 to 2137 nodes.

**Figure 2.**
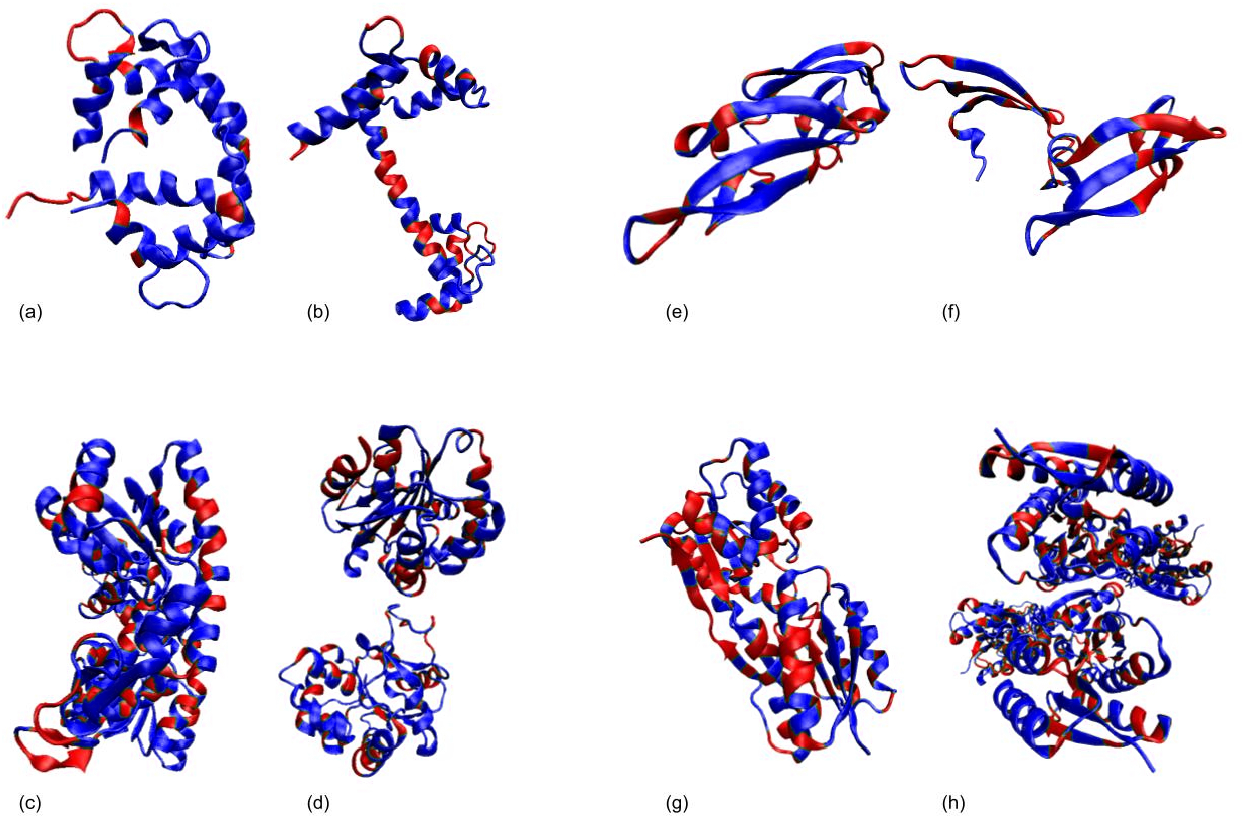
Open and closed conformations of sample proteins. Rigid residues produced by the Kinari software are shown in red, and flexible residues are shown in blue. (**a**) CaM Closed conformation, 1*CTR*. (**b**) CaM Open conformation, 1*CLL*. (**c**) AdK Closed conformation, 4*AKE*. (**d**) AdK Open conformation, 1*AKE*. (**e**) CVN closed conformation, 2*EZM*. (**f**) CVN closed conformation, 15*LE*. (**g**) RBP closed conformation, 2*DRI*. (**h**) RBP open conformation, 1*URP*.

**Figure 3.**
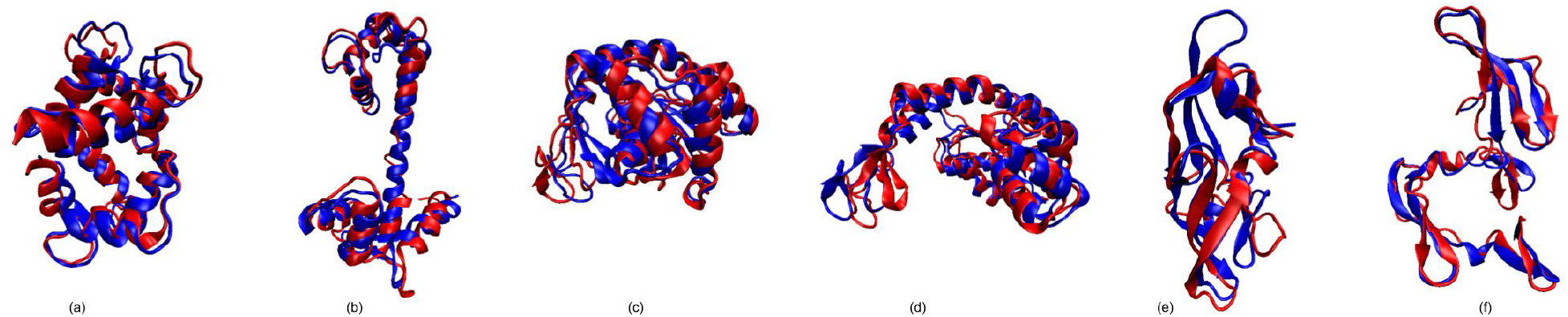
Superimposition of the simulated structure (red) and the original structure (blue). (a) 1*CTR*, (b) 1*CLL*, (c) 1*AKE*, (d) 4*AKE*, (e) 2*EZM*, and (f) 1*L5E*.

**Figure 4.**
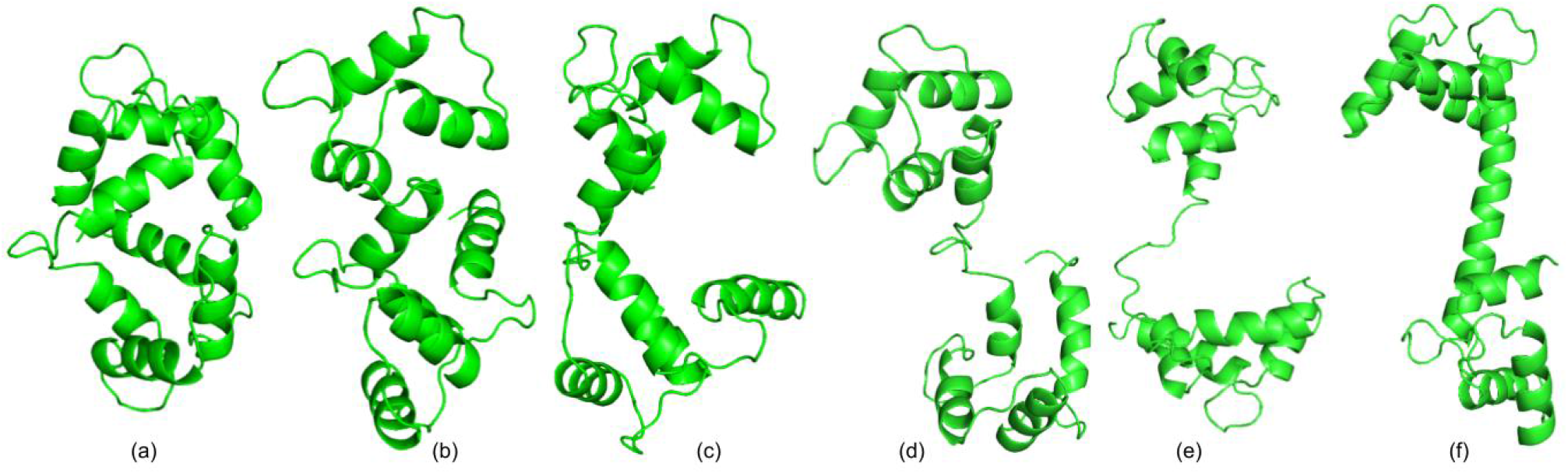
The simulated conformational path for CaM, from 1*CTR*(a) to 1*CLL*(f). Conformations (b) to (e) are the intermediate conformations for this path, taken at approximately equal spaces.

The second pathway that we explored for the Calmodulin protein consisted of the Calcium-free form of Calmodulin (PDB: 1*CFD*) and Calmodulin/IQ domain complex (PDB: 2*F*3*Y*) conformations. Compared to our previous implementation, the algorithm is on average 13 minutes faster for 1*CFD* → 2*F*3*Y*, and 20.87 minutes for 2*F*3*Y* → 1*CFD*. The average tree size changed from 1737 to 1488 for 2*F*3*Y* → 1*CFD*, and from 1432 to 1353 for 1*CFD* → 2*F*3*Y* (Table 2).

#### Adenylate Kinase (AdK)

AdK is a phosphotransferase enzyme (an enzyme that catalyzes phosphorylation reactions), responsible for catalyzing the phosphoryl transfer between two ADP (adenosine diphosphate) molecules to produce ATP (adenosine triphosphate) and AMP (adenosine monophosphate) [41]. The reason AdK is an important molecule when it comes to conformational changes is that it is a signal-transducing protein, and in order to regulate protein activity, there should exist a balance between conformations [42].

*AdK* consists of 214 amino acids and has two small domains, LID and NMP, and a CORE domain [43]. The CORE contains residues 1-29, 68-117, and 161-214, the LID domain in residues 118-167, and the NMP domain in residues 30-67. Research suggests that AdK has two relevant conformations; Open conformation in the unbound structure, 1*AKE*, and a closed conformation 4*AKE*, shown in Figure 2, (d) and (c), respectively. During the conformational transition from the open conformation to the closed one, the most extensive change happens in LID and NMP domains, while the CORE domain stays relatively rigid. The initial lRMSD between the two conformations is 7.14Å. We consider the two pathways for the two conformations of AdK; 1*AKE* → 4*AKE*, and 4*AKE* → 1*AKE*. The average runtime for the 1*AKE* → 4*AKE* path is 228.94 minutes faster than our previous work (Table 1, Refer to section 2.4 for details on runtime analysis). The average run time for 4*AKE* → 1*AKE* is 164.31 minutes faster. Figure 5 shows the progression of lRMSD towards the end conformation. The conformational path created by our current algorithm is smoother than our previous one since we attempt to reduce the randomness by focusing on flexible regions. This helps the algorithm to generate more realistic intermediate conformations and converge faster. Table 2 depicts how integrating rigidity analysis has downsized the RRT tree from having 1526 nodes to 1473 for the 4*AKE* → 1*AKE* path, and from 2398 to 1829 for the 1*AKE* → 4*AKE*. The latter shows a significant amount of memory usage reduction.

**Figure 5.**
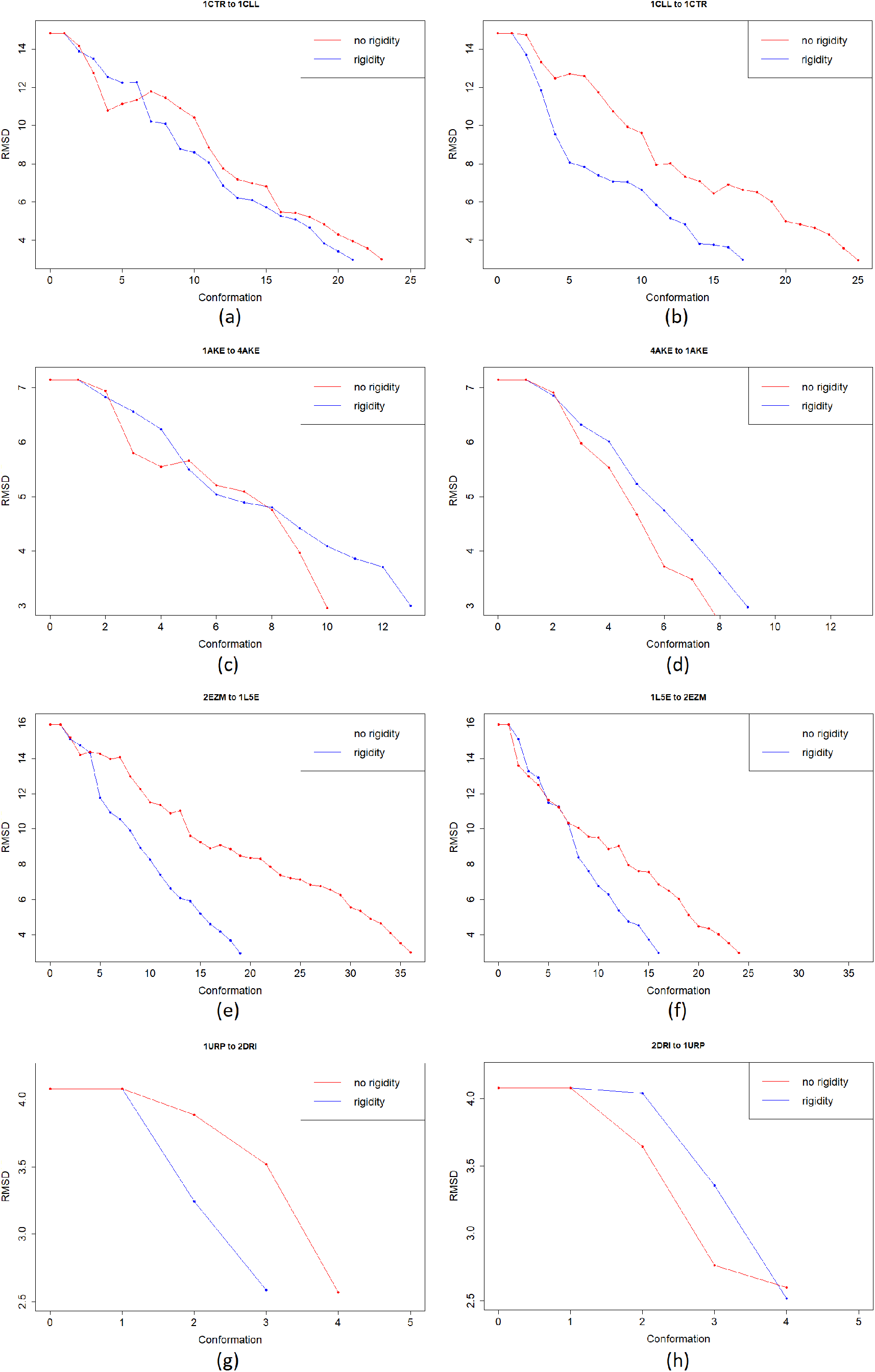
Evolution of lRMSD towards the goal for CaM and AdK. (a) 1*CTR* → 1*CLL*, (b) 1*CLL* → 1*CTR*, (c) 1*AKE* → 4*AKE*, (d) 4*AKE* → 1*AKE*, (e) 2*EZM* → 1*L*5*E*, (f) 1*L*5*E* → 2*EZM*, (g) 1*URP* → 2*DRI*, and (h) 2*DRI* → 1*URP*. The blue line shows the paths obtained by our updated algorithm using rigidity analysis and the red line shows the paths obtained by our previous work based on RRT* algorithm.

#### Cyanovirin-N (CV-N)

Cyanovirin-N (CV-N) is a potent antiviral fusion inhibitor of HIV and many other viruses [44]. It contains 101 amino acids and has two domains with a 30% sequence identity. The domain-swapped dimer has a higher antiviral affinity than the monomer. With a high energy transition barrier, the two forms can be in a solute dissolved into a solvent. Specific mutations can have an effect on the energy barrier and on stabilizing alternative conformations. We attempt to mimic the unpacking of these repeat domains, from the intertwined monomeric conformation to the domain-swapped dimer conformation, for a single chain. 1*L*5*E* is the domain-swapped dimer of CV-N in solution, and 2*EZM* is the solution NMR structure of Cyanovirin-N. The closed and open conformations of CVN are displayed in Figure 2, (e) and (f), respectively, with the rigid parts of protein shown in red. The two conformations have an initial lRMSD of 15.92Å. For the 2*EZM* → 15*LE* path, the updated algorithm runs on average 3.18 minutes faster, and for the 15*LE* → 2*EZM* path it runs 17.5 minutes faster on average (Table 1). The evolution of lRMSD towards the end conformation is shown in Figure 5. Like AdK, the new path is smoother.

#### Ribose Binding Protein (RBP)

In order to function correctly, conformational changes are essential in bacterial periplasmic receptors in chemotaxis and transport. These conformational changes allow ligands to enter and exit and enable correct interactions of ligand-bound proteins. We considered two conformations of the Escherichia coli ribose-binding protein, for which the small backbone moves only happen in the hinge region. The secondary structure elements in the two domains and the amino acids in the binding pocket are relatively rigid.

The open (PDB: 1*URP*) and closed (PDB: 2*DRI*) conformations have dissimilar surfaces in the regions known to be of importance in chemotaxis and transport, that will change their interactions with the membrane components. Conformational changes are necessary for the function of bacterial periplasmic receptors in chemotaxis and transport. Here, we used two conformations of the protein from *Escherichia Coli* (*E.coli*). In *E. coli RBP* (Ribose Binding Protein) small-scale backbone movements are restricted to the hinge region, whereas the secondary structure elements in the two domains and the amino acids in the binding pocket adopt essentially the same conformations [45]. It seems certain that the conformational path that links the forms described here is that followed during ligand retrieval, and in ligand release into the membrane-bound permease system.

The closed and open conformations of RBP are displayed in Figure 2, (g), and (h), respectively, with rigid parts shown in red. For the 2*DRI* → 1*URP* path, the updated algorithm runs on average 21 minutes faster, and for the 1*URP* → 2*DRI* path it runs 18.67 minutes faster on average (Table 1).

## 4. Discussion

In this project, we present and compare different approaches to using rigidity analysis in order to find proteins’ conformational pathways. The use of flexible parts helps us get better results in terms of time, and in most cases, memory usage 2. We examine three scenarios. The first one is the same as our previous work (without using rigidity analysis results). For the second one, we start our algorithm with flexible residues as an input to the algorithm to use instead of all the residues. And in the last scenario, our algorithm not only uses the flexible residues at the beginning but also a few times during the process. Based on the yielded average of runtimes and tree sizes, the third scenario provides us with the best results concerning runtime and memory. By providing the flexible residues to our algorithm, we help decrease randomness and search mostly among suitable candidates instead of randomly chosen protein residues to manipulate. Consequently, this approach yields smoother paths that are less randomized in a faster time. In our future studies, we aim to compare the intermediate conformations and the conformational pathways that our algorithm generates with available experimental results. We intend to investigate how close these pathways are to actual conformational pathways and how we can improve them.

## Author Contributions

All authors contributed equally to the writing. FA and RD conducted the analysis.

## Institutional Review Board Statement

Not Applicable.

## Informed Consent Statement

Not Applicable.

## Data Availability Statement

Not Applicable.

## Acknowledgments

The tests were run on the supercomputing facilities managed by the Research Computing Department at the University of Massachusetts Boston, *Chimera* which is a heterogeneous distributed memory high performance compute cluster (AMD Opteron 6128 processors, 2.0Ghz).

## Abbreviations

The following abbreviations are used in this manuscript:

MDPI: Multidisciplinary Digital Publishing Institute
RRT: Rapidly-Exploring Random Trees
CaM: Calmodulin
AdK: Adenylate Kinase
CVN: Cyanovirin-N
RBP: Ribose-binding Protein
MD: Molecular Dynamics

## Footnotes

1 The EF-hand is a motif that has a helix-loop-helix topology, in which the *Ca*^2+^ ions are coordinated by ligands within the loop, and it is found in a large family of calcium-binding proteins.

